# Sex-Stratified Transcriptomic Meta-Analysis of Alzheimer’s Disease Reveal Brain Region and Sex Specific Dysregulation

**DOI:** 10.1101/2025.06.25.661596

**Authors:** Juan I. Young, Achintya Varma, Lissette Gomez, Michael A. Schmidt, Brian Kunkle, Lily Wang, Eden R. Martin

## Abstract

**Objective:** Alzheimer disease (AD) is a neurodegenerative disorder leading to cognitive decline. Despite growing recognition of sex differences in epidemiology, symptomatology, and clinical outcomes of AD, the molecular mechanisms underlying these variations remain poorly defined. We performed transcriptome association studies of AD aiming to identify sex-specific and sex-dependent transcriptomic profiles that could provide insights into the molecular mechanisms underlying sex differences in AD pathogenesis.

**Methods:** We conducted a meta-analysis of bulk-RNAseq data derived from human postmortem brain studies. Specifically, we analyzed gene expression differences between individuals diagnosed with AD and non-cognitively impaired (NCI) individuals across two key brain regions: the prefrontal cortex and the temporal lobe. We performed stratified differential expression analyses separately in males and females, alongside combined analyses across sexes. Additionally, we assessed the data in relation to known AD genes, proteomic studies, and drug repurposing opportunities.

**Results:** Beyond the genes commonly dysregulated across both sexes, our meta-analyses identified multiple differentially expressed genes (DEGs) between AD and NCI that are either altered in only one sex or show different effects between sexes. Some genes are known AD genes from genetic studies, but others are novel. Correlation with proteomic data suggests that these transcriptional differences have functional significance, potentially contributing to the biological mechanisms underlying sex differences observed in AD. Finally, we identify drug compounds that are potential candidates for treatment.

**Interpretation:** Our findings enhance our understanding of sex-related differences in disease etiology and progression, and underscore the importance of incorporating sex as a critical variable in transcriptomic studies of AD. These insights help pave the way for more precise, personalized medicine approaches that account for sex-specific molecular mechanisms.

## Introduction

Alzheimer’s disease (AD) is the most common cause of dementia in the elderly, characterized by progressive cognitive decline, memory loss, and behavioral changes [1]. Neuropathological hallmarks include extracellular amyloid β (Aβ) plaque accumulation, intracellular neurofibrillary tangles (NFTs) composed of hyperphosphorylated tau, neuronal loss, and reactive gliosis [2]. These changes typically begin in the entorhinal cortex and hippocampus and progressively spread into other cortical areas [3]. AD also associates with altered neurotransmitter signaling, oxidative stress, and neuroinflammation [4–6].

Aging is the primary risk factor for AD, with most cases classified as late-onset AD (LOAD), occurring after 65 years. Incidence increases exponentially with age [7]. Additional factors such as genetic ancestry and genetic risk variants, particularly *APOE*, contribute to disease susceptibility [8]. Another major risk factor for AD is sex. There is growing evidence of sex-related differences in various aspects of the disease, including symptoms, risk factors, and treatment responses [9]. Approximately two-thirds of patients with AD are females [10] and this higher prevalence is not solely due to increased longevity [11]. Recent studies indicate that sex differences in neuropathology, rates of cognitive decline, and neuroimaging persist in age-matched individuals with AD [12, 13]. The APOE-ε4 allele confers a heightened AD risk in females, particularly between ages 65 and 75 [14], and is associated with a stronger accumulation of neurofibrillary tangles in females with mild cognitive impairment [15].

Understanding sex-related molecular mechanisms is crucial for developing effective AD therapies. Sex differences may arise from variations in brain structure, cognitive function, sex hormones influences, inflammatory responses, brain glucose metabolism and differential genetic susceptibility [recently reviewed in 16]. Women exhibit increased levels of total and phosphorylated tau in CSF [17] and tau deposition [18], likely caused by faster tau accumulation due to sex differences in Aβ-dependent secretion [19]. Recent findings on mitochondrial free-carnitine levels suggest metabolic differences also play a role [20]. Additionally, sex disparities in social determinants of health such as education and physical activity may modify AD risk [21,22]

Given the fundamental role of transcriptional programs in brain development, function, and maintenance, unbiased large-scale transcriptome analyses offer valuable insights into sex-related differences in AD pathophysiology and risk. Prior transcriptome studies have identified gene expression alterations in AD, but sex-related transcriptomic dysregulation remains poorly characterized [23].

To explore the role of sex in AD, we performed a meta-analysis of gene-expression data from human postmortem brain studies, focusing on brain regions susceptible to AD pathology. We compared transcriptomic profiles of AD and non-cognitively impaired (NCI) individuals in two key regions: the prefrontal cortex and temporal lobe. Our approach included stratified differential expression analyses in males and females separately, as well as combined analyses across sexes. This enabled the identification of *sex-specific genes* (those affecting only one sex) and *sex-differential genes* (those influencing both sexes but with statistically different effects on AD). We found that, while expression patterns were largely similar between females and males, brains from individuals with AD exhibit a transcriptomic profile with a distinct sex-specific component, providing insights into sex differences in AD etiology and progression.

## Methods

### RNA-seq datasets

We searched public data repositories Gene Expression Omnibus and ArrayExpress, and the literature, for RNAseq transcriptomic datasets containing AD brain samples with sex information. Our selection criteria required candidate datasets to have a minimum of five samples per disease status (>5 AD and >5 NCI). Since our focus is late-onset AD, we required samples to be from individuals older than 65 years of age.

We found four datasets meeting our criteria. Three were from the RNAseq Harmonization study [24–26] available from the AD Knowledge Portal, which consists of RNASeq data from three independent studies: Mayo (syn8466812), Mount Sinai Brain Bank (syn8484987) and ROSMAP (8456629). The fourth dataset was obtained from Gene Expression Omnibus; GSE125583 [27]. Our analyses focused on samples from these datasets from the pre-frontal cortex (PFC) and temporal lobe (TL), key brain regions affected by the disease’s progression and involved in critical cognitive functions.

### Pre-processing of transcriptome data

Unsupervised clustering algorithms were used to verify the consistency of sex labels with their gene expression data. Clustering was performed using values from Y-chromosome probe values, by implementing k-medoids (k=2) [28] clustering algorithm to generate two clusters. The cluster with the higher Y-chromosome values was classified as male, and any sample within this group labelled as female was excluded from further analyses.

The datasets were stratified based on postmortem tissue type. After filtering out genes with low expression and removing mitochondrial genes, we performed surrogate variable analysis (SVA) [29], separately in females and males to obtain covariates that account for hidden batch effects. A variance stabilized count matrix was then used for principal component analysis (PCA) to detect sample outliers, which were subsequently excluded from further analysis.

### Association Testing

We conducted both sex-combined analyses with a full model that includes sex and sex-by-AD interaction, as well as sex-stratified analyses in males and females separately. We focused primarily on identifying gene expression associated with AD diagnosis. In the sex-combined analysis, for each gene, we fit the full linear regression model: *log_2_ (expression) ∼ intercept + AD + age at death + sex + (sex × AD) + sv (n) + (sex × sv (n))*, where *AD* and *sex* are binary terms indicating AD or NCI and male or female, respectively. These main effects are equivalent to the *log_2_ (FC)*, where *FC* is the fold change in expression values between the binary groups. The last terms in the model, *sv (n)*, are the set of *n* surrogate variables estimated by SVA and *(sex × sv (n))* is an interaction that allows for different surrogate variables in females and males. For sex-stratified analyses, the same model was fit to male and female samples separately, excluding the sex and sex-interaction terms. The R program DESeq2 [30] was used for model fitting, providing tests for differential expression and interactions between the sexes (i.e., sex differences).

Meta-analysis was used to combine effect estimates from different studies within each brain region (PFC and TL), requiring results from at least two studies within each brain region. Effect sizes for the main effect of *AD* on gene expression from the sex-stratified analyses, as well as the *sex* term and interaction term for *sex × AD* from the unstratified datasets, were meta-analyzed separately using inverse variance method [31]. We corrected for multiple testing by estimating false discovery rate (FDR).

The magnitude of the estimate of the intercept (i.e., “base-mean” in software) from the models fit in the individual datasets gives a way to assess when read counts may be too low. Specifically, we excluded results from genes with a base-mean less than a given threshold for any of the datasets included in the meta-analyses for either the male or female stratified analyses. Cut-off values (9.0 for TL and 6.1 for PFC) were determined based on the highest Y-chromosome gene expression values observed in female samples, which were considered to be background detection levels. An exception was made for the sex-chromosome genes. Specifically, to capture 17 Y-linked paralogs with low or undetectable expression in females, we applied the base-mean threshold only to male datasets. These Y-chromosome genes were tested in males only but can be compared to results from their X-linked paralog in females.

### Identifying sex-specific genes and sex-differential genes

We classified genes into two groups based on false-discovery rate (*FDR*) and unadjusted *p* values (*p*): *Sex-specific genes* are those significantly associated with AD diagnosis in one sex (*FDR*<0.05) but not in the other sex (*p*>0.05) and showed a nominally significant sex-by-disease interaction (*p*<0.05); and *Sex-differential genes* are those with significant associations with AD diagnosis in both sexes (*p*<0.05) but with nominally significantly differences in effect size between the sexes (*sex × AD* interaction *p*<0.05).

### Pathway enrichment analysis

Pathway enrichment analyses were performed for DEG lists. A user-provided list of 16,727 transcripts expressed in the PFC and 17,181 expressed in the TL were used as the background. 15,411 and 16,563 background transcripts remained in PFC and TL, respectively, as some of the transcripts were not mapped or functionally annotated. Pathways were considered enriched if they met a threshold of *FDR* < 0.05. Metascape was used to perform pathway membership and biological pathway enrichment [32]. P-values were calculated based on the cumulative hypergeometric distribution and *q*-values for *FDR* were calculated based on the Benjamini-Hochberg procedure. The most statistically significant term within a cluster was chosen to represent the cluster. EnrichR [33] was used with the up- and down-regulated gene lists to identify drug repurposing compounds for future work with the tool Drug Perturbations from GEO Down and Up, respectively. We recognize that the enrichment tests provided by Metascape and EnrichR assume independence of genes going into each set, which is not strictly correct for gene expression data. Therefore, we should not interpret p-values/FDR from these tests alone without context. These tools are most useful for generating groups of genes and ranking pathways, and when combined with other supporting evidence can aid interpretation.

## Results

### Datasets

Our meta-analyses included two datasets that used RNA-seq to measure gene expression in brain samples from prefrontal cortex (PFC) and three from the temporal lobe (TL) from four different studies. Specifically, for PFC we used data from the dorsolateral PFC (DLPFC) and the anterior prefrontal cortex (BA10) and for TL we used data from temporal cortex, the fusiform gyrus and the superior temporal gyrus (BA22). The data from the BA10 and BA22 regions both come from the same study (MSBB) with tissues sampled from the same individuals. Table 1 summarizes the distribution of age, sex and AD status for the datasets included in our analyses. After exclusion of samples failing to meet our inclusion criteria and those with discrepancies in recorded sex status versus expression-derived sex status, we had 477 PFC samples and 589 TL samples for analysis.

**Table 1.** Samples used in analyses from non-cognitively impaired (NCI) individuals and individuals with AD by sex with average age [range] in each dataset. DFPLC: dorsolateral prefrontal cortex; BM10: Brodmann area 10; BM22: Brodmann area 22; TC: temporal cortex; FuG: fusiform gyrus.

| Dataset | Brain region | Meta-analysis group | Female |  | Male |  |
| --- | --- | --- | --- | --- | --- | --- |
|  |  |  | Count (N) |  | Count (N) |  |
|  |  |  | Mean Age[age range] |  | Mean Age[age range] |  |
|  |  |  | NCI | AD | NCI | AD |
| ROSMAP | DLPFC | PFC | N=47 | N=108 | N=39 | N=45 |
|  |  |  | 84.74[71.96-90] | 88.51[72.40-90] | 81.96[67.37-9] | 87.75[75.98-90] |
| MSBB | BM10 | PFC | N=40 | N=69 | N=36 | N=40 |
|  |  |  | 85.82[72-90] | 85.38[72-90] | 79.23[66-90] | 80.95[68-90] |
| Total PFC |  |  | N=87 | N=177 | N=75 | N=85 |
| MSBB | BM22 | TL | N=31 | N=55 | N=29 | N=34 |
|  |  |  | 84.5[74-90] | 85.55[72-90] | 79.92[66-90] | 80.88[68-90] |
| MAYO | TC | TL | N=37 | N=49 | N=41 | N=33 |
|  |  |  | 86.69[68-90] | 82.24[66-90] | 84.31[68-90] | 83.67[66-90] |
| GSE125583 | FuG | TL | N=33 | N=91 | N=37 | N=119 |
|  |  |  | 88.05[75-100] | 86.18[66-103] | 86.05[71-100] | 82.61[67-97] |
| Total TL |  |  | N=101 | N=195 | N=107 | N=186 |

### Meta-analysis of AD transcriptomes

Since some of the data (Table 1) were generated from different brain regions within the same individuals (specifically data in MSBB), we performed separate meta-analyses of the PFC and TL data to preserve independence of samples within each analysis. For each brain region, we retained only genes with measurements in more than one dataset, resulting in 16,779 genes in the PFC analysis and 17,784 genes in the TL analysis. After removing mitochondrial genes and genes with low read counts, our final meta-analyses included 16,727 genes in PFC and 17,165 genes in TL.

Our results from association tests between AD and gene expression stratified by sex showed that the AD-associated transcriptomic profiles are quite similar between males and females in both brain regions. The Spearman correlation of effect sizes between sexes across all genes was 0.25 in PFC and 0.64 in TL (Figure 1). Among significant (*FDR*<0.05) differentially expressed genes (DEGs) between AD and controls in both males and females, these correlations increased to 0.97 in both PFC and TL, indicating that genes associated with AD in both sexes tend to have similar effects in males and females. Figure S1 shows volcano plots for the sex-stratified analyses. As expected based on the correlations of effect sizes, we see that the most significant results are usually shared by both females and males, particularly in TL, which has a more balanced ratio of males to females.

**Figure 1.**
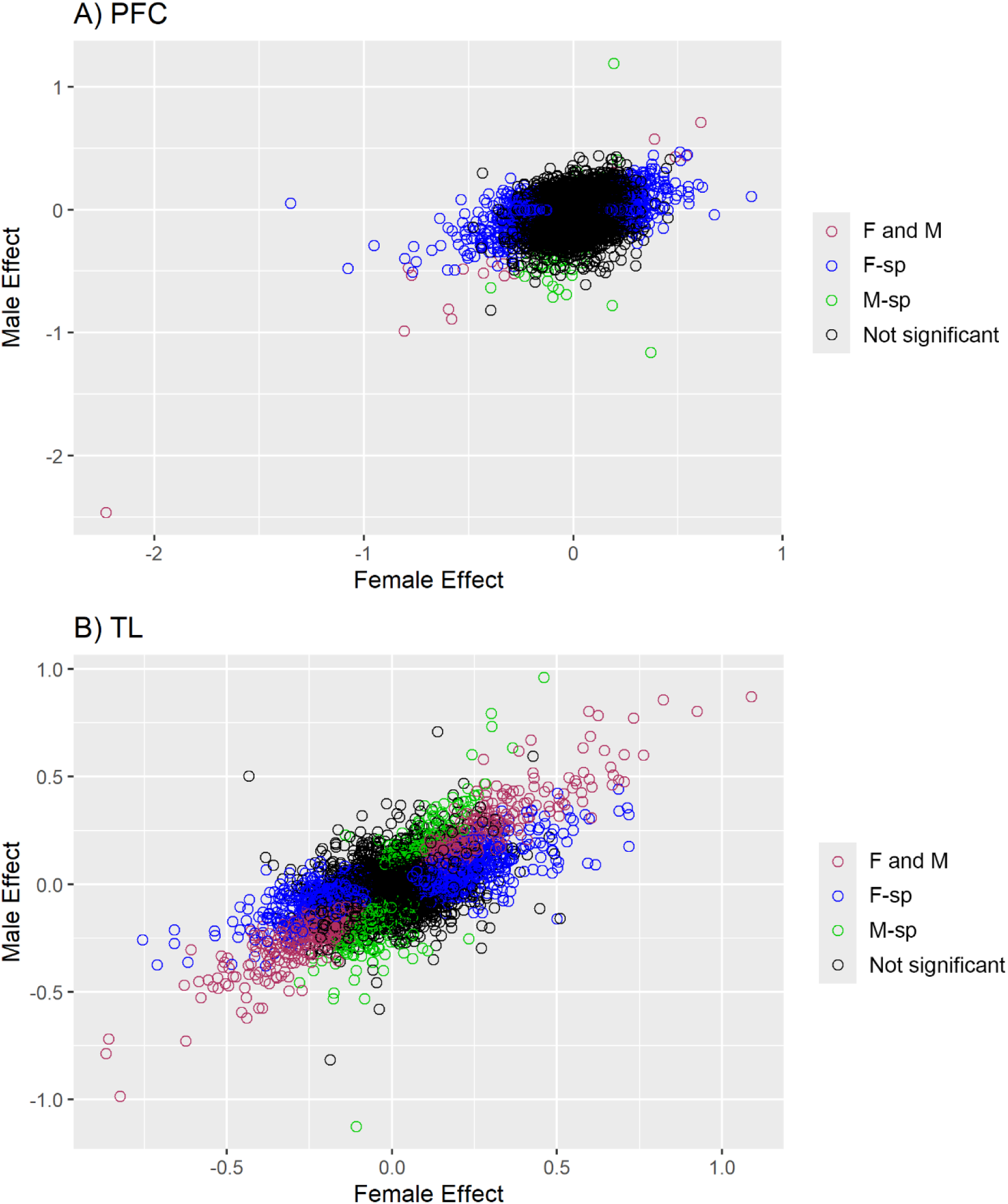
Scatter plots of female versus male effect sizes from the sex-stratified meta-analyses in **A)** PFC and **B)** TF. Results are colored by whether they are significant (*FDR*<0.05) in both females and males (F and M), significant only in the females (F only), significant only in males (M only), or not significant in either females or males. Effect size is the estimate of the coefficient for the main effect in the sex-stratified meta-analyses, which equivalent to log_2_(FC) (FC=fold change in AD versus NCI).

Table S1 lists genes that are FDR-significant (*FDR*<0.05) in both sexes (48 in PFC and 1188 in TL). These DEGs identified in each brain region are significantly enriched with members of the KEGG pathway “hsa05022: Pathways of neurodegeneration - multiple diseases”, compared to the overall meta-analyzed transcriptome (*FDR*=0.03 for both PFC and TL). Pathway enrichment analysis further revealed that genes downregulated in AD in both sexes are primarily involved in synaptic transmission. The most significant KEGG pathway in the PFC is “hsa04726: Serotonergic synapse” (*FDR*=0.008) and in the TL is “hsa04721: Synaptic vesicle cycle” (*FDR*=0.001). Despite the functional overlap, the genes that contributed to the enrichment in synaptic pathways in the PFC and TL are different. On the other hand, the genes upregulated in AD are enriched in chromatin regulation in the PFC, with “hsa03083: Polycomb repressive complex” as the most significant pathway (*FDR*=0.014). In TL, upregulated DEG were enriched most significantly in “hsa04810: Regulation of actin cytoskeleton” (*FDR*=0.0002) and “hsa04510: Focal adhesion” (*FDR*=0.0003). The DEGs belonging to each of the pathways are listed in Table S2.

Seventeen genes that are DEGs in both sexes are shared between the two brain regions (highlighted in green in Table S1). All show a consistent direction of effects in meta-analyses in both tissues: eight genes were downregulated and nine upregulated in both tissues (Figure 2). These 17 shared genes have previously been identified as differentially expressed in additional AD brain regions with the same direction as in this study, as documented in the AGORA database (https://agora.adknowledgeportal.org) (Figure S2).

**Figure 2.**
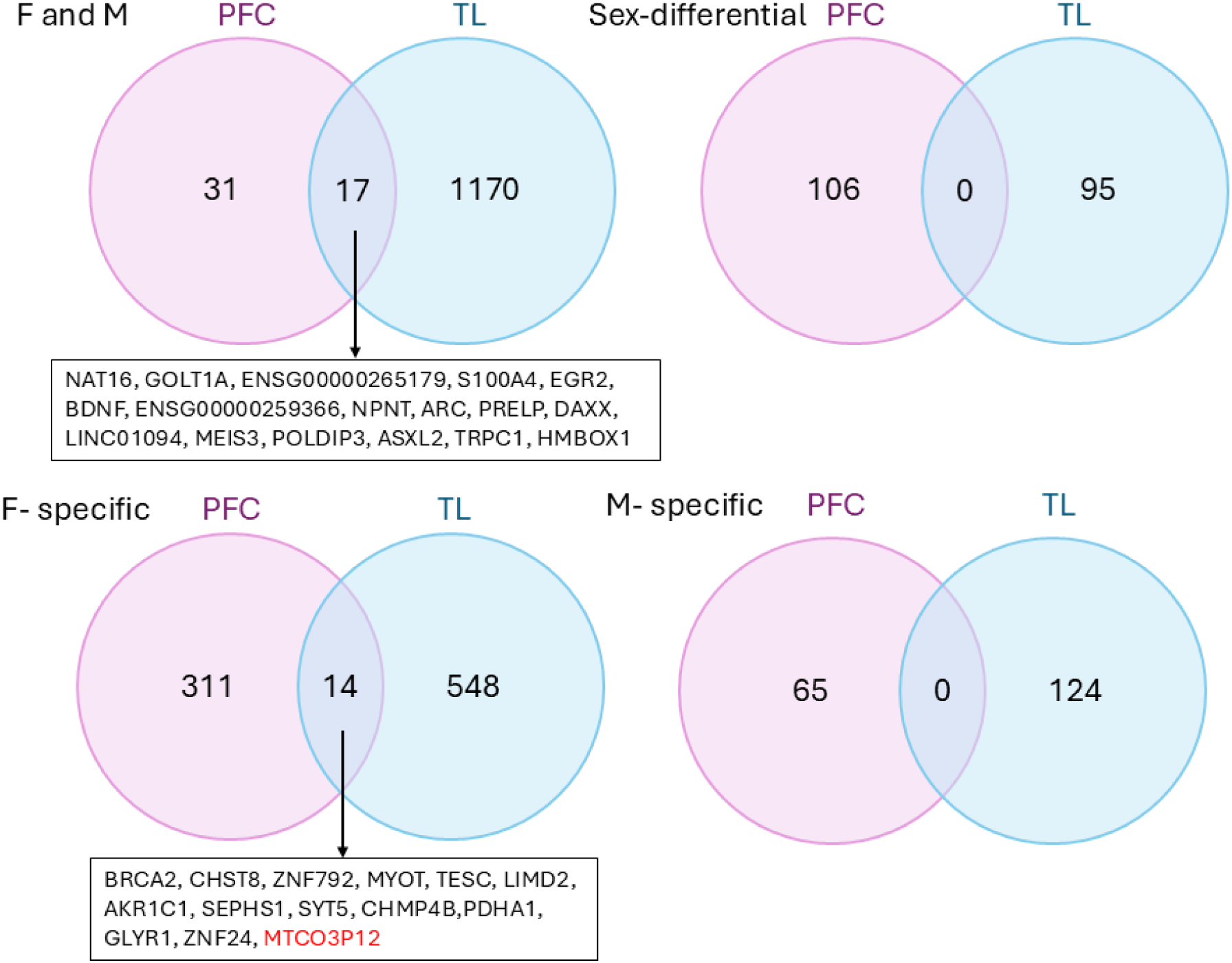
Venn diagrams showing overlap between tissues (PFC and TL) for female-specific genes, male-specific genes and genes significant in both.

Overall, AD-associated gene expression changes are more similar in males and females than they are different (Figure 1); however, in this study we are particularly interested in the differences between the sexes. To capture these differences, we defined two categories of genes: *Sex-specific genes* (*FDR*<0.05 in one sex and *p*>0.05 in the other and with sex-by-disease interaction *p*<0.05) and *Sex-differential genes* (*p*<0.05 in both sexes and sex-by-disease interaction *p*<0.05).

#### Sex-specific Genes

Table 2 presents the number of genes identified as differentially expressed exclusively in one sex, with gene names and statistics provided in Table S3. Among the significant results, there are approximately twice as many genes with negative effect sizes than with positive effect sizes, except for female-specific genes in TL, which have a more balanced distribution of negative and positive effects. This preponderance of negative effects suggests that most sex-specific expression differences in AD are due to down-regulation of gene-expression.

**Table 2.** Counts of sex-specific genes in PFC and TL. We required genes to be present in more than one dataset and passing a base-mean filter in all datasets (base-mean must be >6.1 for PFC and >9 for TL). Negative and positive (-/+) counts refer to the direction of effect size: negative is under-expressed in cases, positive is over-expressed in cases.

| Region | Female Specific (-/+) | Male Specific (-/+) | Total Genes Tested |
| --- | --- | --- | --- |
| PFC | 326 (216/109) | 65 (43/22) | 16744 |
| TL | 562 (264/298) | 124 (85/39) | 17181 |

The top sex-specific genes show concordance with the AGORA database in terms of general direction of effect; however, we identified some significant genes that are not reported as significant in the AGORA dataset. Figure S3 provides examples of sex-specific genes in our data that are not identified as DEGs in the AGORA sex-stratified data. For instance, the AD-GWAS gene *GPC6*, which we identified as a female-specific DEG in the PFC (*FDR*<0.03, *log_2_(FC)*=-0.17) is not reported as a significant DEG in AGORA for females (*p*=0.37) nor males (*p*=0.29). A similar scenario is seen for male-specific DEGs in the PFC, where genes such as *ZNF23*, *HSPA6*, *KCNN4* and *TXNRD2* show no significant evidence of association in AGORA yet are *FDR* significant in our sex-specific analyses. Additionally, *MTCO3P12* and *BRCA2*, which are female-specific DEGs in our analyses of both the PFC and the TL are not reported as significant in either sex within AGORA.

GO term enrichment analyses revealed that female-specific genes downregulated in AD are enriched in biological processes related to synaptic transmission such as the “Modulation of chemical synaptic transmission” (*FDR*=0.0008 and *FDR*=0.0003 for female-specific genes downregulated in PFC and TL, respectively). The set of genes belonging to this pathway in the female-specific DEGs in the two tissues showed limited overlap, with only *SYT5* shared between them. Although female-specific u*p*regulated genes in AD from PFC did not show enrichment in any biological process and only “Protein Modification” (*FDR*=0.003) and “Ribonucleoprotein biogenesis” (*FDR*=0.02) in the TL, we did observe a significant overlap with genes upregulated in in the disease pathway Huntington disease, “Genes Up-Regulated In Human Symptomatic HD BA9 Vs Control GSE129473” (*FDR*=4.1 ×10^-7^ and 2.7×10^-18^ for PFC and TL, respectively). Similarly, the female-specific downregulated genes showed significant overlap with “Genes Down-Regulated In HD Markers In Human Prefrontal Cortex GSE64810” (*FDR*=7.5 x10^-16^ and *FDR*=1.0 x10^-6^ for PFC and TL, respectively). We did not find significant enrichment of any biological processes for male-specific DEGs in either PFC or TL regardless of the direction of change.

#### Sex-differential Genes

Some genes show nominally significant effects in stratified meta-analyses for both sexes (*p*<0.05) and also a significant *sex x AD* interaction (*p*<0.05), indicating different effect sizes between females and males. Our meta-analyses identified 106 such genes in PFC and 95 in TL (Table S4). When applying a stricter threshold requiring FDR-significance in the sex-stratified results (Table 3), this list was reduced to a single gene in PFC and 21 genes in TL. In PFC, the sex-differential gene identified is the *ARC* gene. Relative effects are shown in Figure 3, where you can see that *ARC* is under-expressed in AD cases relative to controls in both males and females (negative slopes for both lines), but with a significantly stronger effect (steeper slope) in females. Interestingly *ARC* is also nominally significantly under-expressed in male versus female controls, as indicated by a significant *sex* effect from the interaction model (*log_2_(FC)*=-0.506, *p*=0.011); however, there is no significant difference in expression between male and female cases, both of which show the lowest levels of expression. Thus in PFC, female controls have the highest level of *ARC* expression, male and female cases have the lowest, and male controls have intermediate expression. Additionally, *ARC* is FDR-significant in both males and females in TL, showing the same trends with females having a stronger negative effect than males (*log_2_(FC)=*-0.423 in females vs. *log_2_(FC)=*-0.282 in males), but with no evidence of *sex x AD* interaction (data not shown).

**Figure 3.**
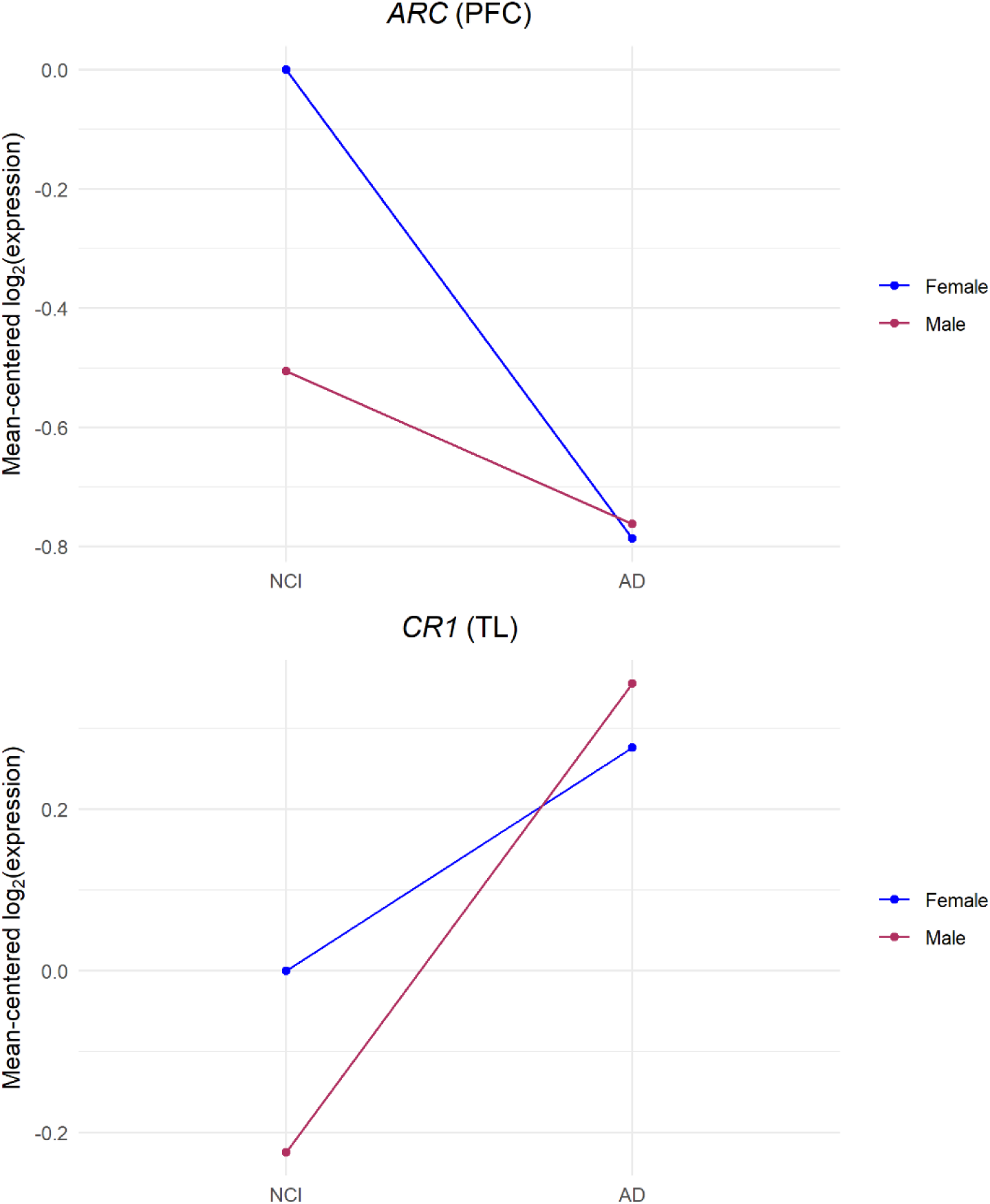
Interaction plots showing the expected values of log_2_(*expression*) for NCI and AD groups in females and males from the combined regression model. log_2_(*expression*) values are centered such that the expected value in NCI females is 0.

**Table 3.** Meta-analysis results from sex-differential DEGs (FDR <0.05 in both sexes and interaction p<0.05) in PFC and TL. INT: *sex x AD* interaction

|  |  |  | FEMALES |  |  | MALES |  |  | INT |  |
| --- | --- | --- | --- | --- | --- | --- | --- | --- | --- | --- |
| Brain Region | Gene | Chr | Effect | FDR | Effect Dir | Effect | FDR | Effect Dir | p | FDR |
| PFC | ARC | 8 | -0.787 | 0.00030 | -- | -0.472 | 0.04215 | -- | 0.041 | 0.610 |
| TL | CPLX1 | 4 | -0.391 | 2.84E-13 | --- | -0.195 | 0.00544 | --- | 0.006 | 0.525 |
|  | SMAD4 | 18 | 0.182 | 1.32E-11 | +++ | 0.104 | 0.00134 | +++ | 0.015 | 0.586 |
|  | NPTXR | 22 | -0.296 | 7.84E-11 | --- | -0.173 | 0.00013 | --- | 0.021 | 0.618 |
|  | TM7SF2 | 11 | -0.257 | 2.32E-10 | --- | -0.115 | 0.02904 | --- | 0.009 | 0.553 |
|  | PLCE1 | 10 | 0.315 | 2.72E-10 | +++ | 0.162 | 0.01051 | +++ | 0.037 | 0.678 |
|  | KCTD11 | 17 | 0.247 | 4.12E-09 | +++ | 0.135 | 0.01003 | ++- | 0.038 | 0.682 |
|  | JPT1 | 17 | -0.197 | 4.16E-09 | --- | -0.095 | 0.00742 | --+ | 0.035 | 0.670 |
|  | STAG1 | 3 | 0.159 | 1.70E-08 | +++ | 0.068 | 0.02630 | +++ | 0.011 | 0.566 |
|  | ARL6IP1 | 16 | -0.149 | 2.47E-08 | --- | -0.080 | 0.01231 | --- | 0.028 | 0.647 |
|  | CGREF1 | 2 | -0.197 | 4.57E-08 | --- | -0.097 | 0.01691 | --- | 0.029 | 0.656 |
|  | ZNF704 | 8 | 0.165 | 2.07E-07 | +++ | 0.086 | 0.01328 | +++ | 0.039 | 0.689 |
|  | PDE5A | 4 | 0.223 | 2.61E-07 | +++ | 0.116 | 0.03153 | ++- | 0.025 | 0.639 |
|  | LCA5 | 6 | 0.194 | 7.68E-07 | +++ | 0.081 | 0.04924 | +++ | 0.030 | 0.656 |
|  | COL26A1 | 7 | -0.260 | 1.00E-06 | --- | -0.133 | 0.03872 | --- | 0.047 | 0.704 |
|  | PCYOX1L | 5 | -0.248 | 2.41E-06 | --- | -0.129 | 0.02872 | --- | 0.041 | 0.691 |
|  | CEP83 | 12 | -0.161 | 2.43E-06 | --- | -0.085 | 0.04171 | --+ | 0.044 | 0.701 |
|  | OLFM1 | 9 | -0.183 | 1.01E-05 | --- | -0.091 | 0.04454 | --- | 0.048 | 0.706 |
|  | PENK | 8 | -0.455 | 0.00191 | --- | -0.597 | 0.00013 | --- | 0.038 | 0.686 |
|  | SMIM18 | 8 | -0.159 | 0.02299 | +-- | -0.258 | 0.00255 | --- | 0.040 | 0.690 |
|  | CR1 | 1 | 0.276 | 0.03671 | +++ | 0.581 | 7.23E-05 | +++ | 0.048 | 0.706 |
|  | LGI2 | 4 | -0.164 | 0.03702 | --- | -0.355 | 1.21E-05 | --- | 0.025 | 0.639 |

In TL, the 21 sex-differential genes (Table 3) all show same direction of effect in males and females. Of these, 17 show significantly stronger effects in females than males, while only 4 genes show significantly stronger effects in males than females. Among the latter is the known AD gene *CR1*, with an effect size of *log_2_(FC)=*0.581 in males, more than double the effect size of *log_2_(FC)=*0.276 in females. Similar to the *ARC* gene discussed above, differences in *CR1* effect size are mainly driven by differences in expression between female and male controls (though not significant for *CR1*) (Figure 3).

#### Sex-specific and Sex-differential AD-associated DEGs show brain region specificity

Very few of the sex-specific and sex-differential DEGs that we identified are shared between the tissues. In the sex-specific analyses, 14 in female-specific genes are found in both tissues but no male-specific genes are shared (Table S5 and Figure 2). Among the 14 female-specific shared genes, all but one (*MTCO3P12*) showed concordant direction in effect in both tissues. None of the genes classified as sex-differential were shared between brain regions, even when we relaxed criteria to include nominally significant p-values.

#### Meta-analysis of genes on the sex chromosomes

Our meta-analyses included genes on the X-chromosome, expressed in both females and males to a variable degree depending on location and X-chromosome inactivation (XCI). Differential expression between females and males in our analyses is captured by a test of the main effect for *sex* from the combined meta-analyses. We found that expression patterns of X-chromosome genes are consistent with their expected XCI status [34]. Specifically, genes that show significantly higher expression in females than males in our data (negative sex main effect, *FDR*<0.05) are known to escape XCI, such as *ZFX* and *KDM6A* (Table S6A). In contrast, pseudo-autosomal region (PAR) genes, such as *GTPBP6* and *CD99*, tend to show higher expression in males than females (Table S6B), consistent with prior reports [35]. Interestingly the gene *MAGEE2* is the only gene not known to lie in the PAR that is significantly over-expressed in males compared to females (in TL, Table S6A).

We found that the average effect size for sex-specific DEGs on the sex chromosomes and autosomes are similar, suggesting that sex differences in AD-associated differential expression is not strongly affected by localization on sex chromosomes. A total of 63 X-chromosome genes are significantly associated with AD in both sexes (Table S1). Only one X-chromosome gene (*SMC1A*) is significant in the PFC (∼2% of the total 48 genes significant in both sexes in PFC), whereas 62 are significant in TL (∼5% of the total 1188 genes significant in both sexes in TL). Several additional X-chromosome genes show sex-specific or sex-differential effects (Table 4, Tables S3-S4).

Among the XCI-escape genes (Table S6A), *PUDP* is significantly associated with AD in males (*log_2_(FC)*=0.19, *FDR*=0.038), with no nominal associations in females in PFC. In contrast, in TL, 9 of the 17 escapee genes (not counting *MAGEE2*) are significantly associated with AD in females (*FDR*<0.05). Among them, *STS* is also FDR-significant in males, with the same direction of effect. Though all of these genes show differences in expression between males and females, none show significant interaction effects between the sexes, suggesting they may be broadly relevant AD and do not show strong sex-specific or sex-differential effects. Looking across tissues, the six XCI genes that we report in PFC are also tested in TL (Table S6A), but only *PUDP* is significantly associated with AD in both brain regions. Interestingly, this gene is significantly upregulated in males in PFC (*log2(FC)*=0.19, *FDR*=0.018) and significantly downregulated in females in TL (*log_2_(FC)*=-0.09, *FDR*=0.038).

Some sex-chromosome genes are known X/Y paralogs, with expression measured separately for their X and Y versions (Table S6C). Our analyses included 35 such genes, consisting of 16 X/Y paralog pairs and the BCOR/BCORL1/BCORP1 trio. The 16 Y-chromosome paralogs are expressed only in males (and some are from a single dataset), but we included them in the analyses to enable comparisons between the expression of their X counterparts in females. In PFC, none of the gene paralogs are FDR-significant for association with AD in males, but three X-chromosome paralogs are FDR-significant in females, *TMSB4X, PRKX and ZFX.* Though not meeting FDR-significance, *PRKX* and the Y-chromosome paralog, *TMSB4Y*, do show nominal significance in males, with similar effect sizes to those observed in females for *PRKX* and *TMSB4X*, respectively. In TL, the same three X-chromosome paralogs, *PRKX*, *TMSB4X* and *ZFX*, are FDR-significant in both sexes, with consistent effect directions across the two sexes. The Y-chromosome paralogs *BCORP1* and *GYG2P1* are FDR-significant, both under expressed in males with AD compared to NCI individuals. The X-chromosome paralogs for *BCOR* and *BCORL1* show nominally significant association in with AD in females in the opposite direction (over expressed) of *BCORP1*. The X-chromosome paralog *GYG2* is not significant in females but is nominally significant in males with an opposite effect direction (over-expressed) of its Y-chromosome paralog *GYGP1*. There are no significant interactions between the sexes for the X-chromosome paralogs. We could only test for significant sex effects for the X-chromosome paralogs; we found that most are significant and all of those have negative sex effects, meaning that expression is higher in male controls than female controls. These findings suggest that, similar to PAR genes, these paralogs may escape complete dosage compensation.

#### Overlap with known AD genes

We assessed the overlap between genes previously identified in AD-GWAS studies [36–39] and the genes identified as DEGs in our meta-analyses. Of the 149 known AD-GWAS genes that we meta-analyzed, 22 and 45 are differentially expressed in at least one of our analyses (female or male stratified analyses, both females and males, sex-specific or sex-stratified analyses) in the PFC and TL, respectively (Table 5, details in Table S7). Twelve of these DEGs are found in both brain regions. If we restrict the overlap analysis to AD-GWAS genes nominated as *bona fide* AD-GWAS candidates by the Alzheimer’s Disease Sequencing Project (ADSP) Gene Verification Committee, then there are 82 AD-GWAS genes included in our meta-analyses. Of these we observed 12 in PFC and 25 in TL that are DEGs in at least one of our analyses. Twelve of these (highlighted in green in Table S7) are DEGs in both brain regions. In the PFC, one of the verified AD-GWAS genes (*SORL1*) exhibits sex-differential expression, being under-expressed in AD females and over-expressed in AD males. Two sex-specific DEGs in the PFC overlapped with AD-GWAS genes (*TRANK1* and *GPC6*). In the TL, 15 verified AD-GWAS genes are DEGs in both sexes, one gene (*CR1*) was identified as sex-differential, and seven genes as sex-specific (*VWA5B2, SPDYE3, ZCWPW1, NYAP1, BCKDK, IL6R and CST3*).

#### Genes containing androgen and estrogen response elements

Sex differences in AD-associated gene expression may be influenced by differential responses to sex hormones. To test whether sex-specific or sex-differential AD genes were enriched for hormone-responsive targets, we assessed the presence of functional estrogen response elements (EREs) and androgen response elements (AREs). To this end, we compared the overlap of genes that we identified as sex-specific and sex-differential with the genes identified as targeted by either estrogen receptor alpha [40] or by androgen receptor [41] (Table S8). In the prefrontal cortex (PFC), 6.3% and 5.4% of our identified DEGs overlap with estrogen- and androgen-responsive targets, respectively. In the TL, the overlap is 5.6% for estrogen and 8.3% for androgen targets. These values are comparable to background levels observed across all expressed genes in each dataset (PFC: 6.2% estrogen, 7.6% androgen; TL: 6.0% estrogen, 7.2% androgen). Further, the proportion of genes in the sex-specific and sex-differential DEGs were not higher that what was observed for DEG in both sexes in either brain region, for which no hormone-specific enrichment is expected. Thus, AD DEGs do not show enrichment for hormone-responsive genes beyond what is expected by chance, suggesting sex hormone signaling does not disproportionately drive the observed transcriptional sex differences in AD.

#### Overlap with the AD proteome

To evaluate the extent to which our findings reflect the brain proteomic profile of AD, we compared them with data from a recent study of the DLPFC proteome in the AD brain [42]. Among the genes we found with expression associated with AD in both males and females, 24% (11 genes) in the PFC and 27.5% (319 genes) in the TL were also identified as AD-associated at the protein level as measured by mass spectrometry. In the PFC, all 11 overlapping genes showed consistent directions between transcriptomic and proteomic changes, and in the TL consistency was seen in 93% of genes (289 transcript-protein pairs) (Figure 4 and Table S9). Notably, two genes that showed differential expression with AD in both sexes and in both tissues, *ARC* (downregulated in AD) and *S100A4* (upregulated in AD), were also identified in the brain proteome study with consistent direction in change between expression and proteome studies. In female-specific AD-associated genes, 26% overlapped with the proteome data in the PFC, with 91% showing consistent direction, and 18% overlapped in the TL, with 76% showing consistency. Among the 14 female-specific genes that are associated with in AD in both tissues, 5 genes (*SYT5*, *CHMP4B*, *PDHA1*, *SEPHS1* and *AKR1C1*) also showed significant changes at the protein levels, with all but *CHMP4B* having consistent directions. For the male-specific genes, we observed 15% overlap with the AD-associated proteins in both PFC and TL, with 78% and 65% showing consistent directionality in PFC and TL, respectively.

**Figure 4.**
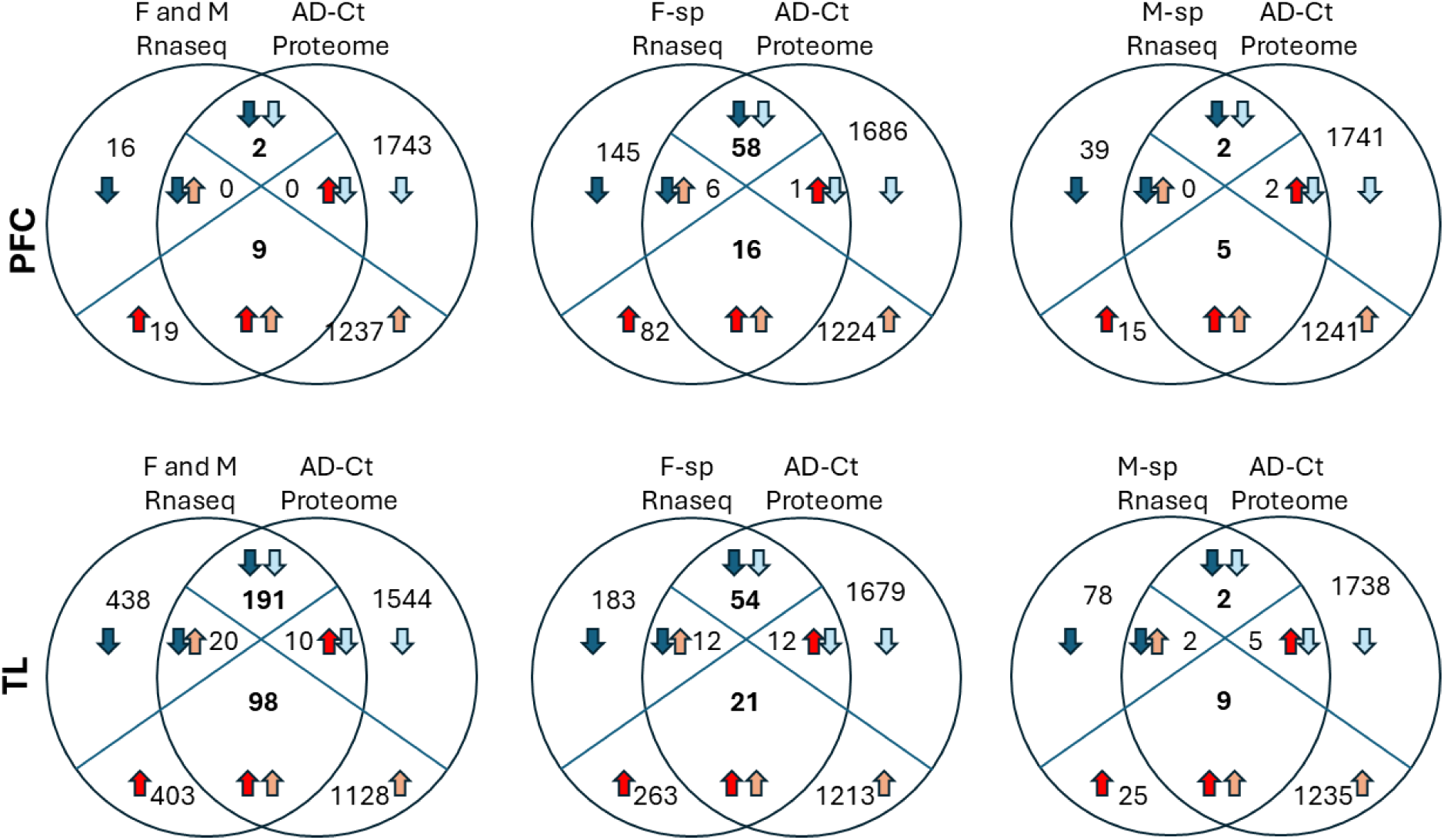
Overlap of meta-analysis results with AD proteome. The proteomic disease-related changes (AD-Ct) were derived from Johnson, E.C.B.,et al. [42]. Blue arrows indicate lower expression in AD (dark blue: transcriptome, light blue: proteome data); red arrows indicate higher expression in AD (dark red: transcriptome, light red: proteome data).

#### Candidate compounds for potential repurposing

To identify candidate therapeutic compounds capable of reversing AD-associated gene expression changes, we queried selected gene sets against drug-induced transcriptional signatures from EnrichR. To prioritize compounds for sex-biased drug repositioning in AD, we restricted the analysis to genes showing consistent directions across all datasets and conducted separate analyses for females and males. In each analysis, we incorporated the DEGs in both sexes, together with sex-differential genes, and sex-specific genes to query the databases, examining both downregulated and upregulated gene sets.

This analysis identified several compounds that are inversely associated with multiple genes in our sex-aware AD gene sets (Table S10A and B). Genes induced by creatine and phenytoin were found to be significantly associated with genes downregulated in females with AD, in both PFC (*FDR*= 0.005 for creatine (GEO dataset GSE5140) and *FDR*=0.007 for phenytoin (GEO dataset GSE2880) and TL (*FDR*=2.72 x 10^-24^ for creatine (GSE5140) and *FDR*= 2.6 x10^-10^ for phenytoin (GSE2880)). These compounds also showed inverse association with genes downregulated in males in the TL (*FDR*= 3.50x10^-5^, GEO dataset GSE5140 for creatine and *FDR*= 6.86x10^-5^, GEO dataset GSE2880 for phenytoin). We also observed that Estradiol signatures were inversely associated with both upregulated and downregulated genes in females with AD (*FDR=*4.75x10^-4^ and 5.85x10^-4^, respectively) and males (*FDR*=0.005 and 3.6x10^-5^, respectively). The aggregate number of genes that show opposite directions of change in the TL AD-associated DEGs vs. the estradiol treated cells is 69 genes (Table S10C). Rosiglitazone emerged as another drug that showed significant inverse associations, showing negative correlations with both upregulated and downregulated genes in females (*FDR*=0.001 and 0.014, respectively) and with genes upregulated in males (*FDR*=1.53x10^-5^).

## Discussion

Despite epidemiological and genetic studies indicating that sex differences exist in AD, our knowledge of the molecular underpinnings of these differences is still limited. Sex differences in AD are understudied; the sex chromosomes have been historically excluded from analyses and sex biases have not been completely considered, likely because of challenges of including sex as a biological variable (i.e. sample sizes). Further, clinical trials are largely biased toward males. In this study, we performed an RNAseq-based transcriptome meta-analyses of postmortem brain samples to obtain a transcriptional sex-aware profile of AD. Hence, in addition to genes dysregulated in AD that are shared by both sexes, we also sought to identify sex-specific and sex-differential gene expression in AD.

Applying a robust analytical pipeline—including sex confirmation, updated gene annotations, use of surrogate variables, and appropriate *sex* x *AD* interaction models—we analyzed data from two AD-relevant brain regions. We identified hundreds of differentially expressed genes (DEGs), either specific to males or females, or shared across both sexes. More DEGs were found in the temporal lobe TL than the PFC, which was expected due to larger sample size and the inclusion of three TL datasets versus two in the PFC. Despite the differences in numbers, there does appear to be brain-region specific gene dysregulation in AD, both for sex-shared and for sex-specific DEGs. In this regard, we detected only 17 genes that had differential expression in both sexes and both brain regions (“core” genes), and only 14 of the female-specific DEGs were shared by the two brain regions, and no male-specific nor sex-differential shared genes.

The set of 17 core DEGs had the same direction of changes in both tissues and both sexes,. Significant expression differences for these genes has also been seen in other brain tissues in other studies, suggesting that their expression is universally affected in AD brains. These expression changes include upregulation of *POLDIP3*, a target of TDP43, a protein intimately linked to neurodegeneration [43]; the proapoptotic factor *DAXX*, involved in amyloid-β toxicity [44]; *S100A4*, shown to have neuroprotective properties [45] and proposed as a plasma biomarker for AD [46]; and *LINC01094*, whose expression has been reported to be associated with age and increased in AD brains [47]. Among the shared downregulated genes are *BDNF*, *ARC* and *EGR2*, implicated in synaptic plasticity [48]. Interestingly, *ARC* and *S100A4* were reported to be also differentially present at the protein level, nominating them as potential therapeutic targets. In fact, both genes have been recently proposed as AD-relevant targets because of their ability to modulate amyloid formation [49,50] and regulate mitochondrial processes [51] and synaptic function [52]. Our finding that *ARC* is also a sex-differential gene, with a more extreme decrease in expression as females go from NCI to AD than in males, suggests that therapies should consider these sex differences and may be sex targeted. A relevant role for *BDNF* in pathogenesis and progression of AD has been repeatedly postulated [53] and altered blood and CNS levels of BDNF and its association with memory and cognitive processes have led to BDNF-based clinical trials for AD [54]. Thus, the most likely outcome of the dysregulation profile of this set of core genes is neuronal dysfunction due to reduced synaptic plasticity and neurotrophic signaling (*ARC*, *EGR2*, *BDNF* and *TRPC1*), neuroinflammation driven by increased astrocyte/microglial activation (*S100A4*, *HMBOX1* and *DAXX)* and oligodendrocyte dysfunction, affecting myelination and neuronal communication (*EGR2*, *PRELP* and *NPNT*). The enrichment of synaptic transmission pathways of genes downregulated across both sexes and brain regions suggests a convergence of affected pathways, even without identical dysregulated genes. This aligns with evidence from in vivo and post-mortem studies showing synaptic loss in AD, which is closely linked to cognitive decline [55,56].

Using the sex-stratified analyses and interaction analysis to ensure significant differences between the sexes, we identified both sex-specific and sex-differential DEGs. We observed repression of female-specific genes participating in synaptic transmission processes in both PFC and TL regions, providing a link between these changes and the suggested accelerated transition from normal cognition to cognitive impairment in females than males [57]. Thus, these data suggest that transcriptomic differences might underly sex-differences in AD risk, onset, progression and severity. Notably, downregulated sex-specific genes, akin to sex-shared genes, demonstrated enrichment in synaptic transmission pathways across both tissues, with minimal overlap observed among the genes contributing to these pathways.

We noted that a few results for sex-specific genes show interesting differences between brain regions. For instance, *MTCO3P12* showed elevated expression in females with AD compared to NCI females in the PFC (effect= 0.85±0.17, *FDR*= 1.83E-04), but had a significant opposite effect in the TL (effect size -0.76±0.16, *FDR*= 6.47E-05). *PDLIM2* also exhibited opposite sex-specific effect directions in the tissues, with under expression only in males in the PFC and over expression only in females in the TL. The behavior of these genes highlights the tissue specificity of sex-dependent gene regulation associated with AD. There are also genes such as *ZNF592*, which we identified as a female-specific gene in the TL, but which exhibits differential expression in both males and females in the PFC. Similarly, *TFRC*, *AGER*, *ENSG00000215014* and *PRSS35* were characterized as male-specific in the PFC but are also significant in both sexes in the TL. In addition, *ZBTB12* is noted as male-specific in the PFC while exhibiting significance in both sexes within the PFC. Finally, *ZFP69B and TMEM109* were identified as a female-specific genes in one brain region and as male-specific in the other. These genes may exhibit sex-dependent expression that is contingent on the particular brain region analyzed; however, it is also possible that they are differentially expressed in both sexes and both brain regions, but are detected in different sexes and regions due to insufficient statistical power.

Because it has been reported that the level of aneuploidy in brains of AD patients is increased compared to normal healthy brains [58–61], there is a possibility that the observed sex-specific DEGs could actually stem from loss of the X chromosome. It has been reported that the frequency of X chromosome aneuploidy in the AD brain is two-fold of what it was observed in NCI females [59]. However, the known mosaic nature and low level (2-5%) of aneuploidy in both unaffected and AD brains suggest that aneuploidy likely does not play a role in sex-specific DEG associated with AD. The subset of X-chromosome genes that we identified as differentially expressed showed both upregulation as well as downregulation in AD, further suggesting that our results are not driven by aneuploidy. Among the X-chromosome DEGs, *PDHA1* is noteworthy because it is identified as female-specific in both tissues with similar negative effect sizes (under expressed in AD). Pyruvate dehydrogenase E1 component subunit alpha (PDHA1) is a pivotal enzyme in glucose metabolism. Deficiency of PDHA1 has been associated with impaired cognitive function in mice [62] and humans [63]. Further, proteomic analysis in AD brains revealed both, decreased protein levels [64] and reduced lysine succinylation [65] of PDHA1 compared to unaffected controls.

Our data suggest that some of the genes identified as bearing AD-associated genetic variants (AD-GWAS genes), also exhibit transcriptional dysregulation, underscoring their relevance for AD pathogenesis and suggesting that variation in gene expression in these genes play an important role in the mechanisms driving disease progression. Interestingly, we observed female-specific dysregulation of two AD-GWAS genes in PFC (TRANK1 and GPC6) and four in TL (ZCWPW1, VWA5B2, SPYDE3, and BCKDK), all of them downregulated. Other notable AD-GWAS genes are *CR1*, which we identified as sex-differential in both tissues and *SORL1*, which was sex-differential in PFC.

Sex differences in transcriptional outputs can arise from multiple contributing factors, including influences of gonadal hormones and sex chromosomes. The results from our meta-analyses, however, did not show a significant enrichment for sex-specific genes on the X chromosome or for genes containing sex hormone responsive elements, suggesting the sex-specific transcriptome in these two brain areas might depend on additional sex-dependent factors such as genetic architecture, gene-by-environment interactions and cellular liability. Among the X-chromosome genes upregulated in AD is *PRKX*, a gene that was identified as upregulated in AD brain oligodendrocytes [66] and encodes a serine threonine protein kinase with a role in neural development.

We focused on identifying compounds capable of reversing gene expression changes associated with AD through the principle of signature reversion: compounds predicted to downregulate genes upregulated in AD and upregulate those that are downregulated. Interestingly, this approach identified estradiol as a drug with significant potential to correct transcriptome changes in the TL. Estradiol is a sex hormone that has been previously shown to downregulate inflammatory cytokines in the central nervous system (CNS) and has neurotherapeutic effects on animal models of AD. Estrogen plays a role in amyloid precursor protein (APP) processing and overall neuronal health by regulating various factors such as BDNF, calcium signaling, excitotoxicity and apolipoprotein E (ApoE) [67]. Further, the prevalence of AD increases in postmenopausal women, suggesting a potential association between lower estradiol levels and disease onset. However, results with hormone replacement therapy in postmenopausal AD women have been inconsistent [68]. Our data suggest that estradiol responses vary between different areas of the brain and therefore local effects should be considered to understand the association between estrogen and AD. We also identified rosiglitazone, a PPARγ agonist used in type 2 diabetes [69], as potentially reversing AD transcriptomic signatures in the TL. While preclinical studies show cognitive benefits via improved insulin signaling and reduced inflammation [70] clinical trials have not supported its efficacy in AD, possibly due to poor CNS penetration or patient variability [71]. Lastly, creatine, a key regulator of bioenergetics, emerged as another candidate. It supports mitochondrial function and ATP regeneration, both of which are disrupted in AD. Recent studies have proposed creatine as a neuroprotective agent with potential for AD prevention and treatment. [72].

In summary, our sex-aware meta-analysis highlights that, beyond the genes commonly dysregulated across both sexes, there also exist a substantial number of genes that exhibit sex-specific patterns of expression. Interestingly, correlation with proteomic data suggests functional significance of these changes. Thus, these distinct gene expression profiles likely contribute to the biological mechanisms underlying sex differences observed in AD. Our data support the utility of transcriptome-based drug repurposing approaches and highlight candidate compounds for further experimental validation in sex- and region-specific models of AD. These findings emphasize the importance of incorporating sex as a critical biological variable in genetic and transcriptomic studies of AD, as well as in the development of targeted therapeutic strategies.

## Acknowledgements

This research was supported by US National Institutes of Health grants R01AG062634 (E.R.M, B.K, L.W.) and U01AG072579 (J.I.Y). The authors acknowledge the assistance of Quinn “The results published here are in whole or in part based on data obtained from the AD Knowledge Portal (https://adknowledgeportal.org). Data generation was supported by the following NIH grants: P30AG10161, P30AG72975, R01AG15819, R01AG17917, R01AG036836, U01AG46152, U01AG61356, U01AG046139, P50 AG016574, R01 AG032990, U01AG046139, R01AG018023, U01AG006576, U01AG006786, R01AG025711, R01AG017216, R01AG003949, R01NS080820, U24NS072026, P30AG19610, U01AG046170, RF1AG057440, and U24AG061340, and the Cure PSP, Mayo and Michael J Fox foundations, Arizona Department of Health Services and the Arizona Biomedical Research Commission. We thank the participants of the Religious Order Study and Memory and Aging projects for the generous donation, the Sun Health Research Institute Brain and Body Donation Program, the Mayo Clinic Brain Bank, and the Mount Sinai/JJ Peters VA Medical Center NIH Brain and Tissue Repository. Data and analysis contributing investigators include Nilüfer Ertekin-Taner, Steven Younkin (Mayo Clinic, Jacksonville, FL), Todd Golde (University of Florida), Nathan Price (Institute for Systems Biology), David Bennett, Christopher Gaiteri (Rush University), Philip De Jager (Columbia University), Bin Zhang, Eric Schadt, Michelle Ehrlich, Vahram Haroutunian, Sam Gandy (Icahn School of Medicine at Mount Sinai), Koichi Iijima (National Center for Geriatrics and Gerontology, Japan), Scott Noggle (New York Stem Cell Foundation), Lara Mangravite (Sage Bionetworks).”

## Author Contributions

J.I.Y, A.V, L.G, B.W.K, L.W and E.R.M contributed to the conception and design of the study; J.I.Y, A.V, L.G,M.A.S, B.W.K, L.W and E.R.M contributed to the acquisition and analysis of data; J.I.Y, A.V, L.W and E.R.M contributed to drafting the text or preparing the figures.

---

**Figure S1.**
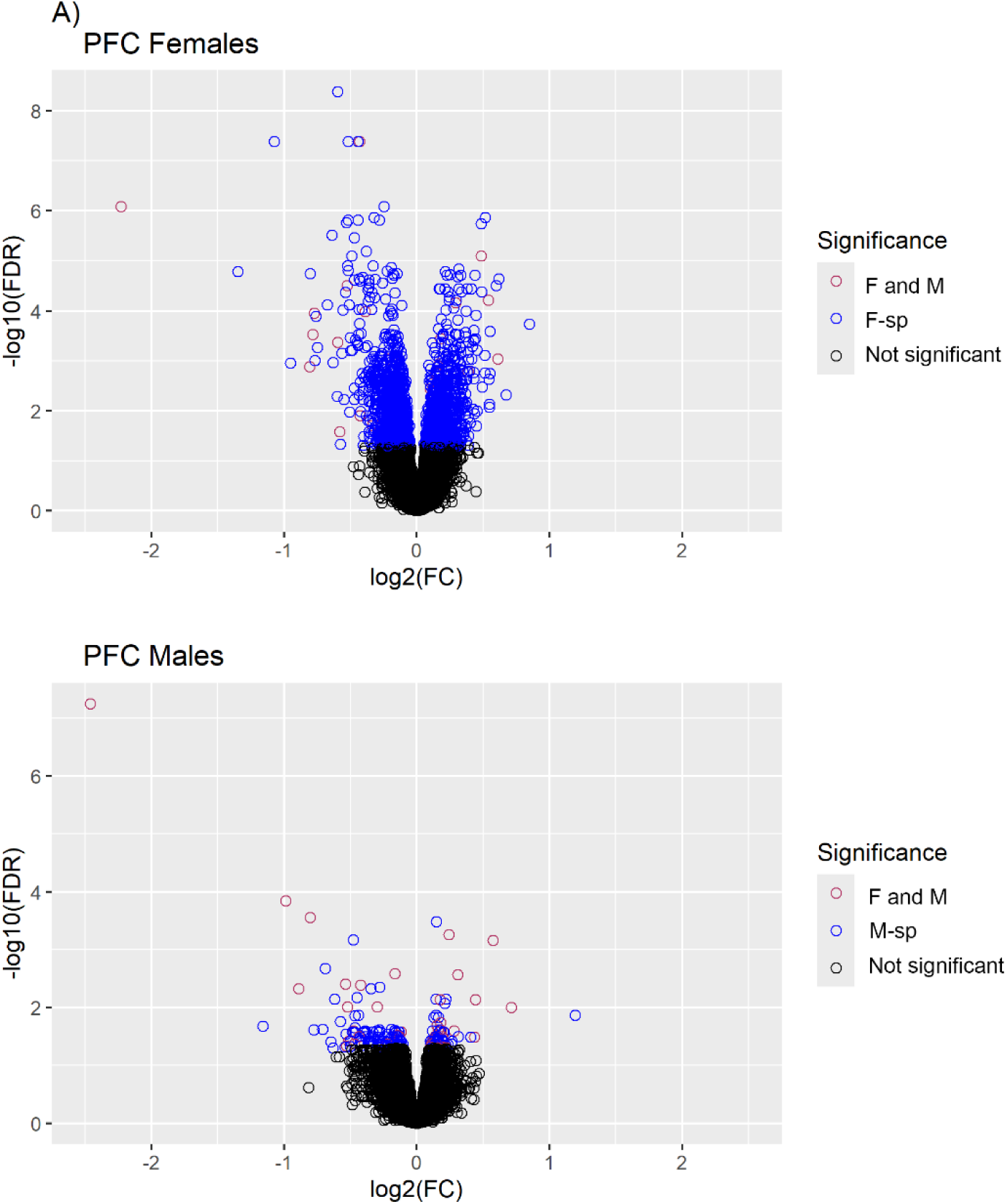
Volcano plots of meta-analysis results in **A)** PFC and **B)** TL from sex-stratified analyses. Results are colored by those that are significant (*FDR*<0.05) in both females and males (F and M), significant only in the females (F only) for the female strata or significant only in males (M only) for the male strata, or not significant in the respective strata. Effect size is the estimate of the coefficient for the main effect from the sex-stratified meta-analyses, which equivalent to *log_2_(FC)* (FC=fold change in AD versus NCI).

**Figure S2:**
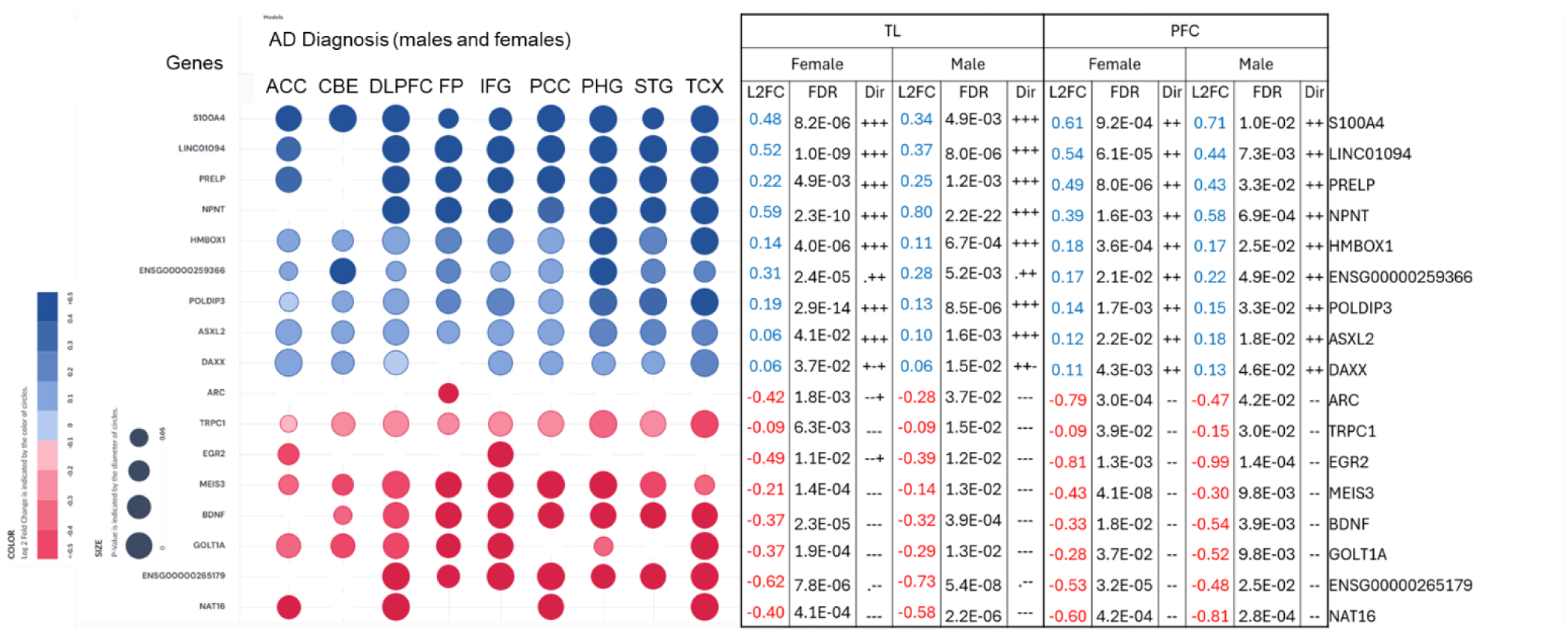
Differential expression in AD of the 17 genes shared by females and males in both the PFC and the TL, as presented in the AGORA website (left) and our metanalyses data (right).

**Figure S3.**
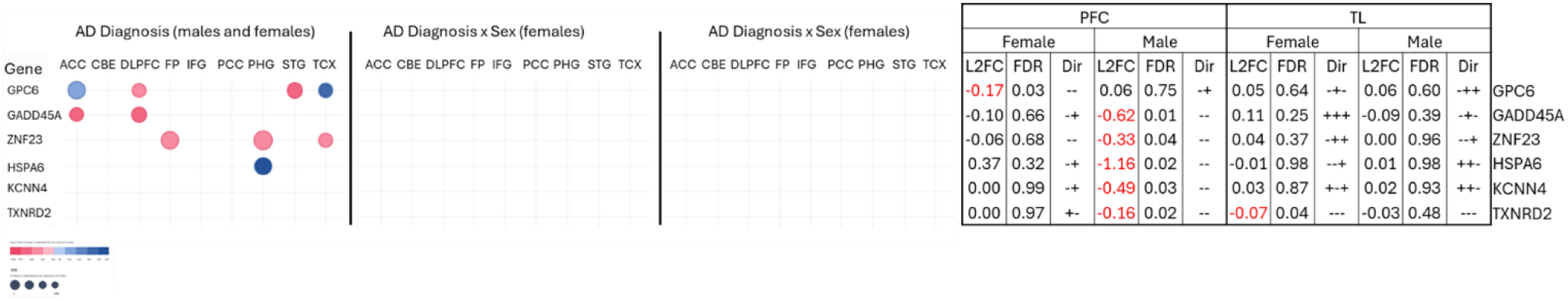
Sex-specific genes in our data, as compared to the AGORA website data. These genes are significant in our study but not identified as significant in the AGORA sex-stratified analysis. *KCNN4* and *TXNRD2* are not identified as DEG by AGORA.

